# RNA polymerases display collaborative and antagonistic group behaviors over long distances through DNA supercoiling

**DOI:** 10.1101/433698

**Authors:** Sangjin Kim, Bruno Beltran, Irnov Irnov, Christine Jacobs-Wagner

## Abstract

Transcription by RNA polymerases (RNAPs) is essential for cellular life. Genes are often transcribed by multiple RNAPs. While the properties of individual RNAPs are well appreciated, it remains less explored whether group behaviors can emerge from co-transcribing RNAPs under most physiological levels of gene expression. Here, we provide evidence in *Escherichia coli* that well-separated RNAPs can exhibit collaborative and antagonistic group dynamics. Co-transcribing RNAPs translocate faster than a single RNAP, but the density of RNAPs has no significant effect on their average speed. When a promoter is inactivated, RNAPs that are far downstream from the promoter slow down and experience premature dissociation, but only in the presence of other co-transcribing RNAPs. These group behaviors depend on transcription-induced DNA supercoiling, which can also mediate inhibitory dynamics between RNAPs from neighboring divergent genes. Our findings suggest that transcription on topologically-constrained DNA, a norm across organisms, can provide an intrinsic mechanism for modulating the speed and processivity of RNAPs over long distances according to the promoter’s on/off state.

## INTRODUCTION

RNA polymerases (RNAPs) carry out the first step of gene expression by transcribing DNA into RNA. Inside cells, a gene is often transcribed by multiple RNAPs. Therefore, it is important to understand not only how a single RNAP transcribes a gene, but how multiple RNAPs transcribe a gene together. Do co-transcribing RNAPs translocate faster (or slower) or dissociate less (or more) frequently than a solo RNAP? If so, what is the mechanism underlying the emergence of the group behavior?

Experiments have shown that when an RNAP runs into a stalled RNAP (arrested by a roadblock or a sequence-specific pause site) it can effectively ‘push’ the paused RNAP (Epshtein and Nudler, 2003; Epshtein et al., 2003; Jin et al., 2010; Saeki and Svejstrup, 2009). This ‘RNAP push’ occurs because a trailing RNAP can prevent a paused RNAP from backtracking or help shift the equilibrium of a backtracked RNAP towards translocation. Since RNAPs often pause temporarily (Landick, 2006), the ‘RNAP push’ effect can increase the apparent transcription elongation rate by reducing pause duration. This model proposes that the rate of transcription elongation increases with the density of RNAPs on the DNA template and therefore with the rate of transcription initiation due to additive ‘RNAP push’ effects (Epshtein and Nudler, 2003; Epshtein et al., 2003). This local cooperation between RNAPs is thought to be most effective for genes with very strong promoters (Epshtein and Nudler, 2003; Proshkin et al., 2010; Saeki and Svejstrup, 2009), such as ribosomal genes, where elongating RNAPs are close to each other due to frequent back-to-back loading onto the DNA (Voulgaris et al., 1999).

The evidence for the elongation rate increasing with the initiation rate through cumulated ‘RNA pushes’ primarily stems from observations made using a promoter (T7 A1) whose strength approaches that of maximally induced ribosomal promoters (Deuschle et al., 1986). Comparatively, the vast majority of genes across cell types have much weaker promoters (see Figure S1 for *Escherichia coli*) (Bon et al., 2006; Pelechano et al., 2010; Schwanhäusser et al., 2011; Taniguchi et al., 2010). The density of RNAPs on the DNA can also greatly vary from gene to gene (Figure S1) (Larson et al., 2014; Mayer et al., 2015; Min et al., 2011; Mokry et al., 2012; Mooney et al., 2009; Pelechano et al., 2009; Vijayan et al., 2011; Wade and Struhl, 2004), implying that RNAPs can be separated by a wide range of distances during transcription elongation. Under these physiological contexts, it remains unknown whether RNAPs traveling at a distance affect each other and therefore show group behavior. It is generally assumed, without concrete experimental evidence, that well-separated RNAPs transcribe a gene the same way as a single RNAP transcribes a gene by itself.

In this study, we examine whether co-transcribing RNAPs can display group behaviors under transcription initiation rates commonly found among *E. coli* genes. We provide evidence that under a wide range of physiological gene expression levels, the rate of transcription elongation does not change with the rate of transcription initiation, suggesting that the ‘RNAP push’ mechanism has a negligible effect on overall RNAP speed under these conditions. However, transcription elongation efficiency of already transcribing RNAPs becomes compromised when the loading of new RNAPs stops due to promoter inactivation. This occurs independent of how active the promoter was before being turned off, as long as there were more than one RNAP on the DNA template. These contrasting results are reconciled by a mechanism in which RNAPs affect each other over long distances, either positively or negatively, through transcription-induced DNA supercoiling.

## RESULTS AND DISCUSSION

### Large changes in transcription initiation rate do not affect the transcription elongation rate

To examine how a modulation of the transcription initiation rate may affect the transcription elongation rate, we used the *lac* operon of *E. coli*, a paradigm of bacterial gene regulation. The activity of the native *lac* promoter can easily be tuned by varying the concentrations of the membrane-permeable inducer isopropyl β-D-1-thiogalactopyranoside (IPTG) (Monod, 1956). This, in effect, modulates the initiation rate and thus the density of, and the spacing between, co-transcribing RNAPs on the DNA. In addition, the lac promoter can be rapidly shut off by the addition of glucose or orthonitrophenyl-β-D-fucoside (ONPF) (Adesnik and Levinthal, 1970). The first gene of the lac operon encodes LacZ, a β-galactosidase whose production can be monitored using the Miller assay (Miller, 1972). Since translation is coupled to transcription in bacteria (i.e., the first ribosome follows the RNAP) (Figure S2) (Kohler et al., 2017; Landick et al., 1985; Miller et al., 1970; Proshkin et al., 2010), the apparent rate of transcription elongation, *r*, can be estimated by dividing the length of the *lacZ* transcript (3,072 nt) by the time span between IPTG addition and the rise in β-galactosidase activity (Jin et al., 1992; Kepes, 1969; Schleif et al., 1973).

For our Miller assay experiments, we used 0.2 or 1 mM IPTG for maximal promoter activity and 0.1 and 0.05 mM for intermediate and low activities, respectively (Figure 1A). Based on genome-wide RNAP profiling (Larson et al., 2014) and reported initiation rates for the *lac* promoter (So et al., 2011), these promoter activities cover a range of RNAP densities commonly observed among well-expressed *E. coli* genes that have important functions in cell physiology (Figure S1). In the ‘RNAP push’ model, *r* increases with RNAP density and hence promoter activity through cumulated ‘RNAP pushes’ (Epshtein and Nudler, 2003; Epshtein et al., 2003). Inconsistent with this expectation, we found that the first functional LacZ enzymes appear at about the same time under high, intermediate and low IPTG concentrations (intercept with the baseline in Figures 1B and S3). In other words, *r* was similar under all tested promoter activities (Figure 1C), despite up to ~4-fold reduction in LacZ synthesis (Figure 1A).

**Figure 1.**
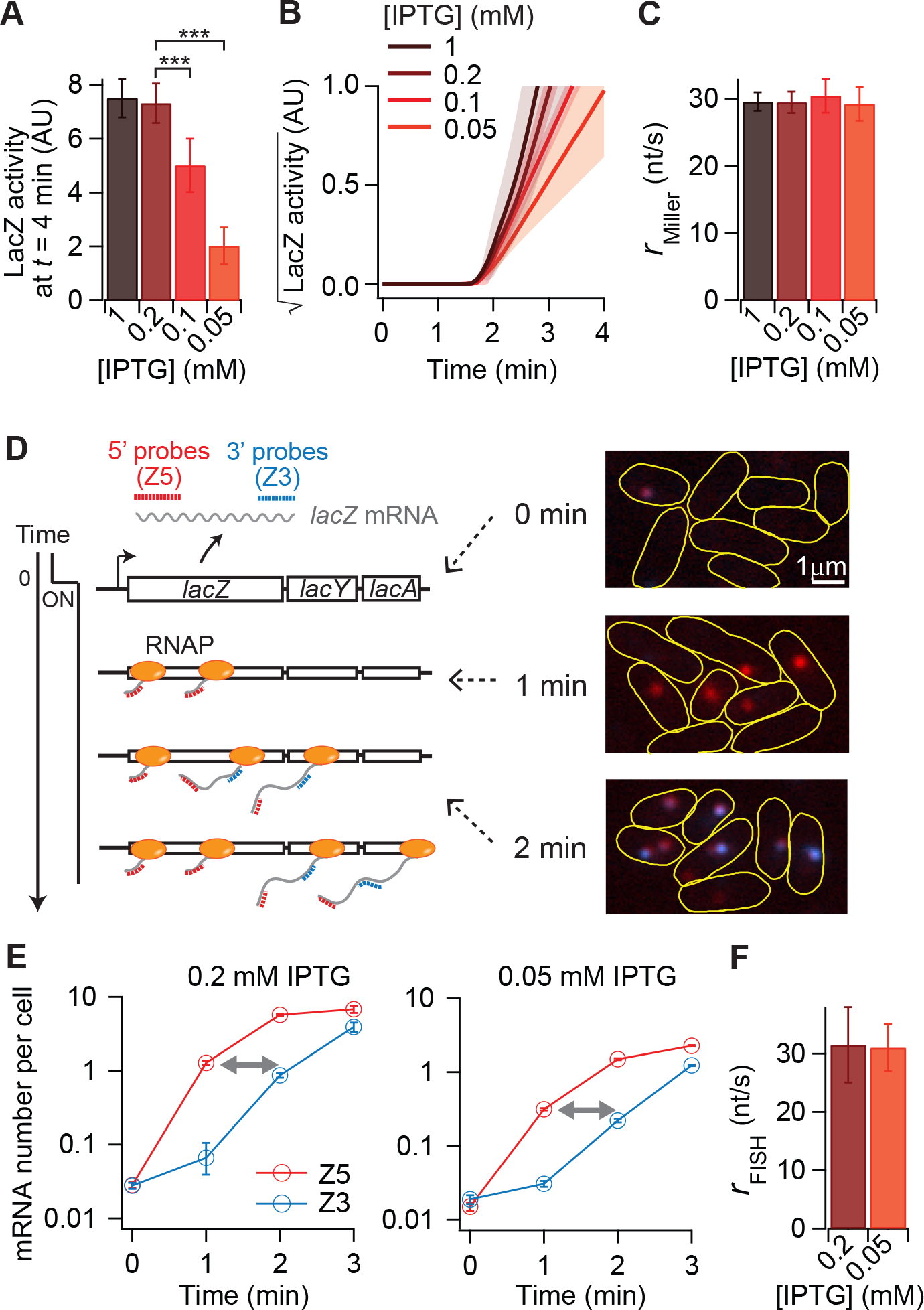
Effect of promoter strength on the rate of transcription elongation. Expression of *lacZ* in wild-type *E. coli* MG1655 cells grown at 30?C was assayed over time by Miller assay (A-C) or single-molecule mRNA FISH microscopy (D-F) following induction with the indicated IPTG concentrations. (A) LacZ activity (after baseline subtraction) measured 4 min after IPTG addition. The three asterisks denote a statistically significant decrease (*P* < 0.001, two-sample *t* test). Error bars show the standard deviations for at least four experiments. (B) Kinetics of the square root of LacZ activity following IPTG addition. The square root was used because the LacZ amount is expected to increase as a function of *t*^2^ (Schleif et al., 1973). Lines and shaded areas indicate the means and standard deviations of two-line fits (a baseline fit from *t* = 0 to the appearance of LacZ and a linear fit of the initial increase in LacZ activity) done on each time-course experiment (example traces are shown in Figure S3). A total of six, eight, six, and thirteen experiments were performed for 1, 0.2, 0.1, and 0.05 mM IPTG conditions, respectively. (C) Apparent transcription elongation rate of *lacZ* at indicated IPTG concentrations. Error bars show the standard deviations of at least three experiments. (D) (Left) Schematic of single-molecule two-color FISH microscopy used to measure lacZ *mRNA* levels over time. Red and blue dotted lines indicate Cy5 or Cy3B fluorescently-labeled oligonucleotide probes that hybridize to 1-kb-long 5’ and 3’ *lacZ* mRNA regions, or Z5 and Z3, respectively. (Right) Overlay of two fluorescence images with pseudo-coloring for Cy5 (red) and Cy3B (blue) at indicated time points after IPTG addition. Data shown at *t* = 0 correspond to that of a sample collected before IPTG addition. (E) Z5 and Z3 numbers per cell over time after IPTG addition. Arrows qualitatively show the time shift in Z3 appearance. Error bars are bootstrapped standard errors of the mean. At least 1200 cells were analyzed per time point. (F) Effect of different promoter activities on the apparent transcription elongation rate of *lacZ*, calculated by dividing the distance between the two probe regions (2000 nt) by the time shift between the Z5 and Z3 mRNA signals. Error bars are standard deviations of five and eight experiments for the 0.2 and 0.05 mM IPTG conditions, respectively. See also Figures S1, S2, and S3.

We verified these results with an independent and more direct method by probing mRNA synthesis over time using two-color single-molecule fluorescence in situ hybridization (FISH) microscopy (Iyer et al., 2016). In this assay, 1-kb regions at the 5’ and 3’ ends of the *lacZ* mRNA (Z5 and Z3, respectively) were visualized at one-minute intervals using different fluorescently labeled probes (Figure 1D). This method provides population-averaged kinetics of transcription elongation based on measurements from thousands of cells. The shift in time between the rise in Z5 and Z3 signals (Figure 1E) represents the time required for the first RNAPs to translocate from the 5’ to the 3’ probe regions and provides another means for calculating the apparent elongation rate (Iyer et al., 2016). Using this approach, we found that *r* was identical under maximal (0.2 mM) and low (0.05 mM) IPTG induction conditions (Figure 1F), in good quantitative agreement with the Miller assay data (Figure 1C).

Our results indicate that modulating the rate of transcription initiation by several folds does not affect the rate of transcription elongation. Under conditions of maximal induction, the *lacZ* gene has an RNAP density that is lower than that of ribosomal genes, but higher than that of most other *E. coli* genes (Figure S1). Thus, the RNAP density produced by the fully induced *lac* promoter is already too low to produce a cumulated ‘RNAP push’ effect large enough to significantly alter the apparent rate of elongation.

### Turning off an active promoter results in apparent slowdown of transcribing RNAPs

The lack of correlation between initiation and elongation rates under common levels of gene expression feeds into the general assumption that well-separated RNAPs do not affect each other’s motion. If this assumption is true, turning off an active promoter—a common natural occurrence when the environment changes—should not have any effect on the apparent elongation rate of already-loaded RNAPs. To our surprise, this is not what we observed. Since our assays report on the transcription elongation rate of the first loaded RNAPs after IPTG induction, we shut off the promoter before the first RNAPs reached the end of the *lacZ* gene by adding an anti-inducer, ONPF or glucose, 90 s after induction with 0.2 or 0.05 mM IPTG. At both IPTG concentrations, LacZ synthesis was significantly delayed following promoter inactivation (Figures 2A and S4). Under these conditions, we detected functional LacZ only at *t* ≈ 160 s, compared to *t* ≈ 110 s when the promoter remained active, indicating an overall decrease in *r* (Figure 2B). Since the conditions were the same for the first 90 s, these results imply that it took about three times longer (160 s – 90 s = 70 s vs. 110 s – 90 s = 20 s) for the first RNAPs to complete *lacZ* transcription following promoter inactivation. This result is remarkable because the first RNAPs were over 2 kb away from the promoter (based on their average elongation rate) when the promoter was turned off, indicating that the ON/OFF state of the promoter has a long-distance effect on transcribing RNAPs. This long-distance effect was not associated with the particularities of ONPF or glucose inhibition, as a decrease in *r* was also observed when the promoter was turned off with rifampicin (Figure S5A).

**Figure 2.**
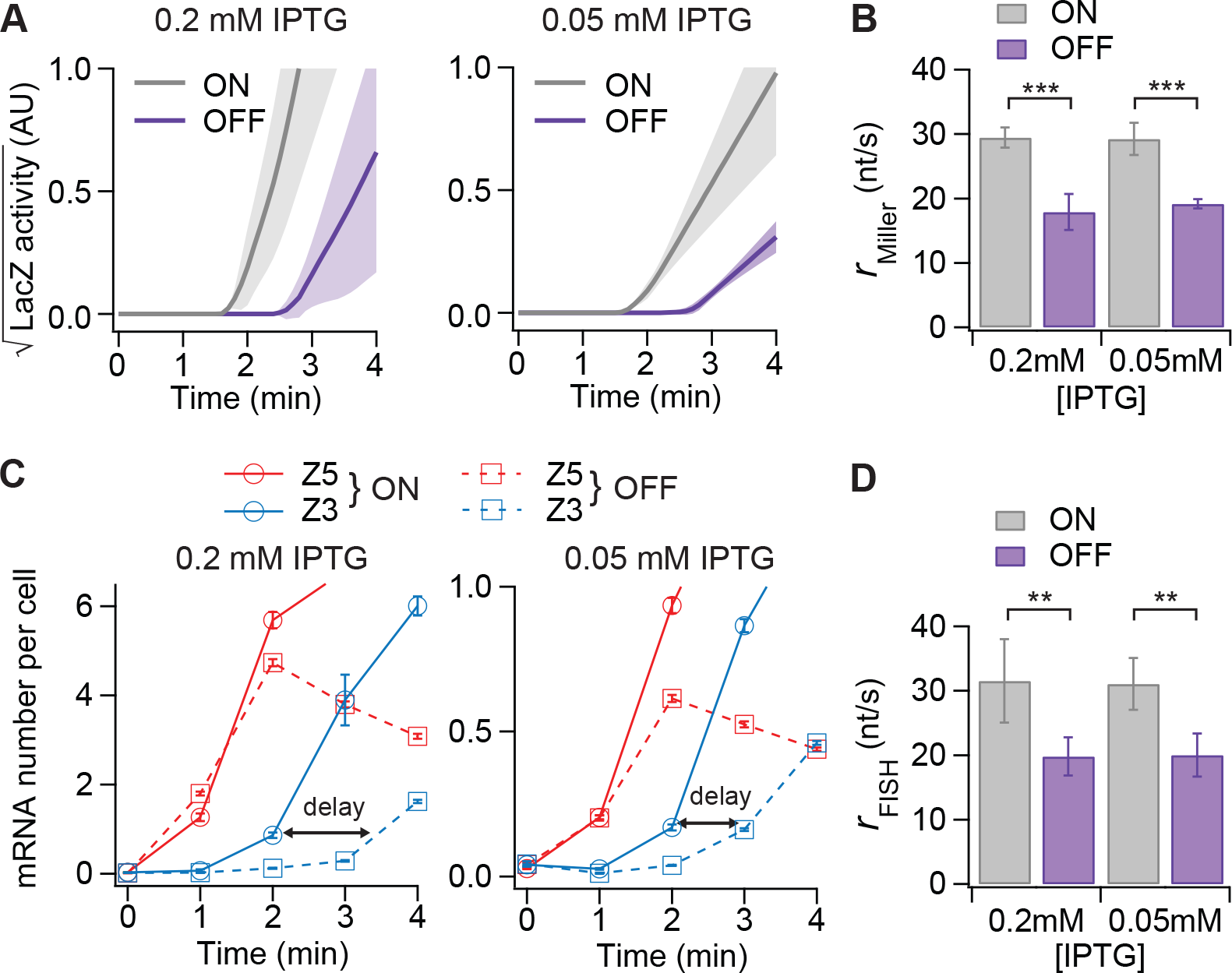
Effect of promoter inactivation on transcription elongation rate. (A) Miller assay results showing the kinetics of the square root of LacZ activity depending on whether the promoter remains induced (ON) or is turned off (OFF). The promoter was inactivated by addition of 5 mM ONPF or 500 mM glucose at *t* = 90 s after induction with 0.05 or 0.2 mM IPTG, respectively. AU, arbitrary units. Lines and shaded areas indicate the means and standard deviations of two-line fits on each time-course trace (*n* = 8 (ON) and 6 (OFF) experiments for 0.2 mM IPTG condition and *n* = 13 (ON) and 11 (OFF) experiments for the 0.05 mM IPTG condition). (B) Effect of promoter inactivation on r measured by Miller assay (as in Figure 1C). *** indicates *P* < 0.001 (two-sample *t* test). Error bars show the standard deviations of replicates described in (A). (C) Z5 and Z3 mRNA numbers per cell over time in FISH microscopy experiments in which the promoter was turned off (OFF) or not (ON) by addition of 500 mM glucose at *t* = 90 s. Black arrows indicate the delay in Z3 appearance from the basal level in the OFF case relative to the ON case. Over 1200 cells were analyzed per time point. Error bars are bootstrapped standard errors of the mean. (D) Effect of promoter inactivation on *r* measured by two-color mRNA FISH microscopy as in Figure 2C. Error bars are standard deviations of *n* = 5 (ON) and 6 (OFF) experiments for the 0.2 mM IPTG condition and *n* = 8 (ON) and 15 (OFF) experiments for the 0.05 mM IPTG condition. ** indicates *P* < 0.01 (two-sample *t* test). See also Figures S4, S5, S6, S7, and S8.

Could promoter inactivation somehow cause the formation of a long-lived pause near the end of the *lacZ* gene? If it did, shutting off the promoter earlier, such as at *t* = 45 s instead of 90 s, would result in the same delay, as the RNAPs should only experience this pause when they reach that pause site near the end of the gene. If, instead, the apparent RNAP slowdown is not linked to the formation of a specific pause, but occurs immediately or shortly after promoter inactivation, turning off the promoter earlier should further delay the first appearance of LacZ activity. We observed the latter (Figure S6), arguing against the formation of a specific pause site and arguing in favor of an apparent slowdown of RNAPs immediately after the promoter is turned off.

We confirmed the long-distance effect of the promoter shut-off on transcription elongation using FISH microscopy experiments in which the promoter either remained on or was turned off with glucose 90 s after addition of 0.2 mM IPTG. The Z5 mRNA signal appeared at the 1-min time point in both cases (Figure 2C). The same timing was expected, as it occurred before glucose addition. However, the first appearance of the Z3 mRNA signal was delayed from the 2-min time point to the 3-min time point when the promoter was shut off compared to when it remained active (Figure 2C). This delay reflects a reduction in *r* (Figure 2D), in agreement with the Miller assay results (Figure 2B). We obtained similar results with 0.05 mM IPTG (Figures 2C, 2D and S5B), indicating that the observed decrease in apparent elongation rate is insensitive to a large change in the density of RNAPs loaded onto the DNA template.

These observations were recapitulated in a Δ*lacYA* strain (Figure S7), thereby ruling out any potential effect from the expression of downstream genes *lacY* and *lacA* (e.g., LacY-dependent positive feedback on transcription initiation (Novick and Weiner, 1957; Ozbudak et al., 2004)). The delay in *lacZ* transcription upon promoter inactivation was also independent of the genomic context, as it was reproduced in a strain in which the *lac* operon is expressed from a plasmid instead of its native chromosomal locus (Figure S8).

### The apparent RNAP slowdown in response to promoter inactivation occurs in vitro with the minimal set of components needed for transcription

To examine whether our promoter shut-off observations are linked to an inherent property of transcription (i.e., independent of other cellular processes), we turned to an in vitro transcription assay. For this, we used a plasmid containing the original lac operon sequence with a two-base mutation in the promoter (*lac*UV5), which is commonly used in in vitro studies because it does not require an activator protein (CAP) for full promoter activity (Noel and Reznikoff, 2000). Since transcription is independent of IPTG in vitro (no LacI repressor), expression from the *lac*UV5 promoter was induced by adding purified *E. coli* RNAPs to the reactions. We found that shutting off the promoter with rifampicin before the first RNAPs completed *lacZ* transcription significantly reduced their apparent speed in vitro (Figure 3A), despite the absence of ribosomes or other cellular factors apart from RNAPs and the plasmid template. This suggests that the reduced efficiency of transcription elongation observed in vivo results from an intrinsic property of transcription.

**Figure 3.**
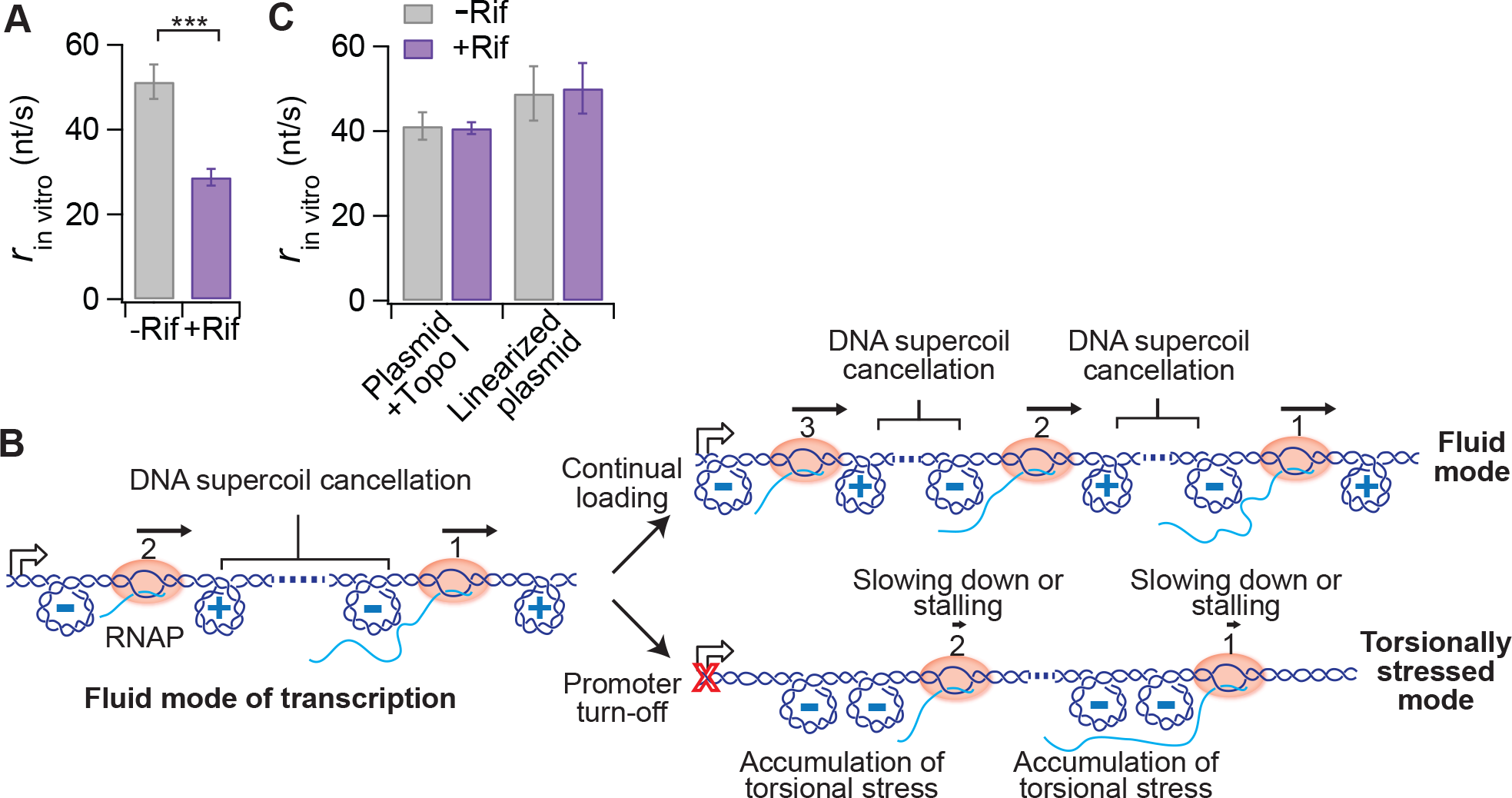
Effect of DNA supercoiling on *lacZ* transcription kinetics depending on the promoter’s ON/OFF state. (A) Apparent transcription elongation rate of *lacZ* measured in vitro using a plasmid containing *lacZYA* driven by the *lac*UV5 promoter. At *t* = 0, purified *E. coli* RNAP holoenzyme was added to induce multi-round transcription. At *t* = 30 s, rifampicin (+Rif) was added or not (-Rif). Error bars are standard deviations of nine (-rif) and seven (+rif) experiments. *** indicates *P* < 0.001 (two-sample *t* test). (B) Schematic showing the proposed model for transcription-driven DNA supercoiling affecting RNAP kinetics depending on whether the promoter remains active or is turned off. See text for details. (C) Same as (A) except in the presence of Topo I or using the linearized plasmid as a template. Error bars are standard deviations of four experiments for each condition.

### Transcription-induced DNA supercoiling mediates two modes of transcription elongation depending on the promoter’s ON/OFF state

How can shutting off a promoter rapidly affect the translocation of RNAPs that are so far away from the promoter? We hypothesized that the apparent slowdown of transcription elongation after promoter inactivation may be related to DNA supercoiling intrinsically generated by RNAPs as they transcribe a topologically constrained DNA template (i.e., a template that cannot rotate). During transcription, individual RNAPs generate negative DNA supercoiling upstream while creating positive DNA supercoiling downstream (Liu and Wang, 1987). On the other hand, it has been shown that accumulation of either negative DNA supercoils upstream (Ma et al., 2013) or positive DNA supercoils downstream of an RNAP (Chong et al., 2014; Ma et al., 2013; Rovinskiy et al., 2012) inhibits the translocation of this polymerase. We reasoned that when two RNAPs transcribe on a DNA template, negative and positive DNA supercoils between RNAPs may cancel out (Figure 3B), as previously hypothesized (Guptasarma, 1996; Liu and Wang, 1987). Therefore, we envisioned that DNA supercoil cancellation by neighboring RNAPs would reduce torsional stress, promoting a more ‘fluid’ mode of transcription elongation (Figure 3B, left). Cancellation requires both positive and negative DNA supercoils to be produced by RNAP translocation, suggesting that RNAP motion is important. In other words, the motion of an RNAP would help that of the next RNAP. DNA supercoil cancellation would also occur between distantly-spaced polymerases because DNA supercoils can quickly diffuse over long distances (van Loenhout et al., 2012). In this context, sustained loading of RNAPs would be important as it would ensure that the level of negative DNA supercoiling behind the last-loaded RNAP (i.e., the one closest to the promoter) does not accumulate beyond an inhibitory threshold (Figure 3B, top).

Such a ‘fluid’ mode of transcription elongation would be abrogated when the loading of new RNAPs stops (i.e., when the promoter is turned off). Accumulation of negative DNA supercoils behind the last-loaded RNAP would cause it to slow down or stall. This slower RNAP would then generate fewer positive DNA supercoils downstream, reducing its long-distance assistance on the translocation of the nearest downstream RNAP through DNA supercoil cancellation. A slowdown or stalling of this downstream RNAP would then have the same negative effect on the translocation of the next RNAP, and so forth. As a result, the disruptive torsional effect on the translocation of the last-loaded RNAP would rapidly propagate to RNAPs far downstream, creating a ‘torsionally stressed’ mode of elongation (Figure 3B, bottom). Under this mode, the slowdown of an RNAP would promote the slowdown of other RNAPs on the DNA, meaning that RNAPs negatively impact each other when the promoter is turned off.

Consistent with our hypothesis, adding type I topoisomerase (Topo I) to the in vitro transcription reaction to remove negative DNA supercoils resulted in similar average elongation rates regardless of whether the promoter remained active or was turned off by rifampicin (Figure 3C). We note that the elongation rate with the constitutively active promoter (no rifampicin) was lower in the presence of Topo I than in its absence (Figure 3C vs. Figure 3A). One possible explanation is that Topo I not only removes the accumulated negative DNA supercoils behind the last RNAP when the promoter is turned off, but also removes negative DNA supercoils in-between RNAPs before they can cancel out with positive DNA supercoils generated by the nearby RNAP. An accumulation of positive DNA supercoils also creates torsional stress that impacts RNAP translocation (Chong et al., 2014; Ma et al., 2013; Rovinskiy et al., 2012), explaining the lower *r* value in the presence of Topo I. To circumvent this problem and prevent accumulation of any type of DNA supercoils, we linearized the plasmid, thereby allowing its free rotation during transcription elongation. Indeed, linearization of the DNA template restored the higher rate of transcription elongation as well as abrogated any effect that turning off the promoter had on the elongation rate (Figure 3C). These results indicate that DNA supercoiling coordinates the change in elongation dynamics according to the ON/OFF state of the promoter.

Altogether, our results support a model in which co-transcribing RNAPs aid each other’s translocation over a long distance through DNA supercoiling cancellation as *long* as the promoter continues to supply new RNAPs onto the gene. This positive interaction between RNAPs over long distances is not cumulative in that it is independent of RNAP density as long as positive and negative DNA supercoils between RNAPs can diffuse toward each other and cancel out. This ‘fluid’ mode of transcription elongation would explain why the apparent rate of transcription elongation on *lacZ* is the same at maximal (0.2 mM IPTG) and low (0.05 mM) levels of induction (Figure 1). Based on RNAP density comparison (Figure S1), most well-expressed *E. coli* genes, including those involved in critical aspects of cellular physiology, are expected to experience a ‘fluid’ mode of transcription elongation as well.

### A solo RNAP displays a slower apparent speed than multiple co-transcribing RNAPs and is not affected by promoter inactivation

According to our model, if there is only a single RNAP per template, as expected for repressed or weakly expressed genes, the absence of torsional stress relief from co-transcribing RNAPs through DNA supercoiling cancellation should result in a reduced transcription elongation rate. This is, indeed, what we observed in Miller and FISH experiments when *lacZ* expression was induced with only 0.02 mM IPTG (Figures 4A-4D). Under this very low induction condition, only a single RNAP is present on the *lacZ* template, based on the observation that the number of Z5 mRNAs per fluorescent spot does not increase over time following IPTG induction, unlike at higher IPTG concentrations (Figure 4E).

**Figure 4.**
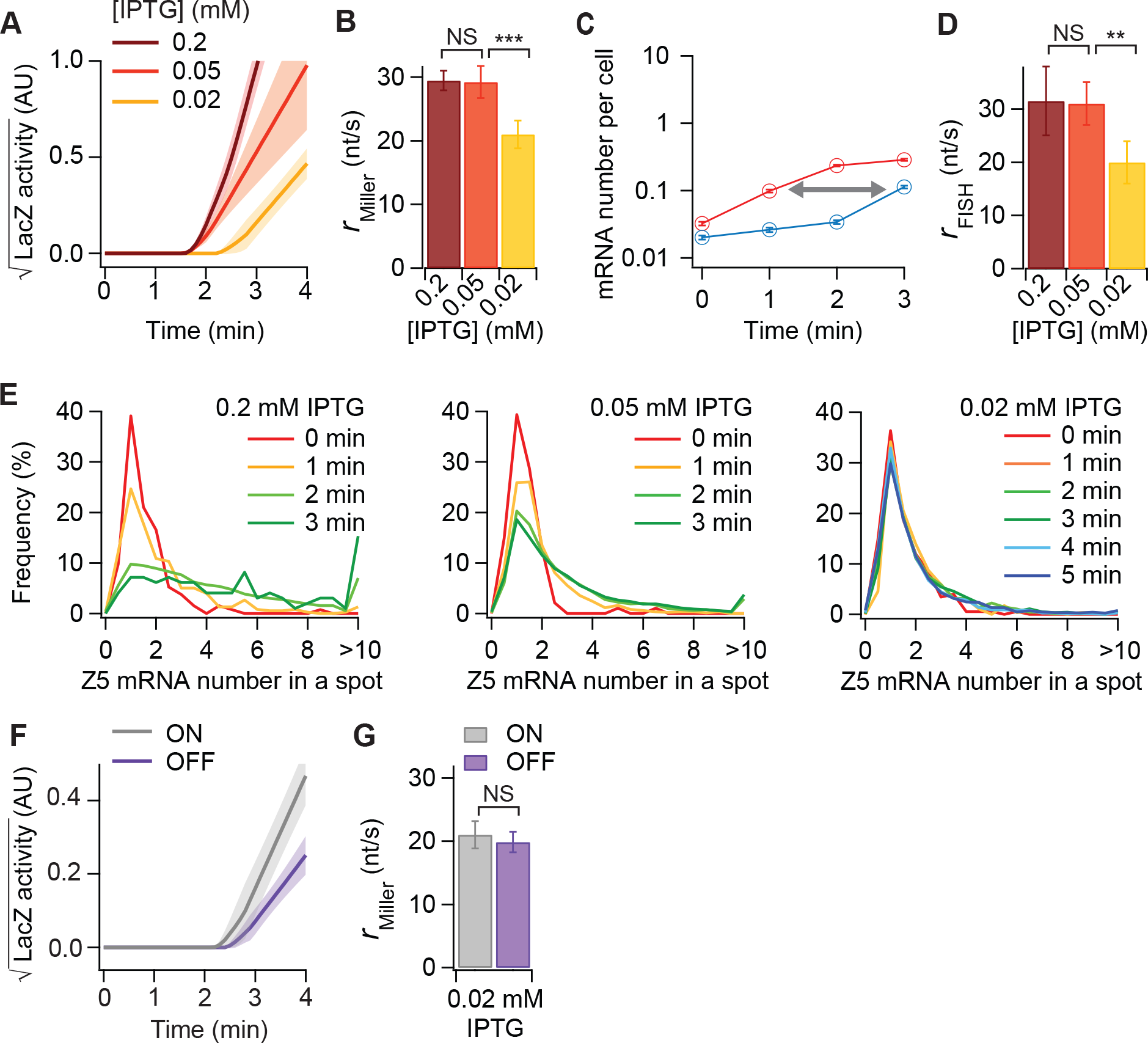
Transcription elongation kinetics when the lac promoter is minimally induced. (A) Kinetics of the square root of LacZ activity following IPTG addition. AU, arbitrary units. Lines and shaded areas indicate the means and standard deviations of two-line fits on each time-course trace from at least three experiments. (B) Apparent transcription elongation rate of *lacZ* at indicated IPTG concentrations. Error bars show the standard deviations of at least three experiments. *** indicates *P* < 0.001 (two-sample *t* test). NS indicates a non-significant difference. (C) Z5 and Z3 mRNA numbers per cell over time after 0.02 mM IPTG addition. The arrow qualitatively shows the time shift in Z3 appearance. Error bars are bootstrapped standard errors of the mean. At least 7000 cells were analyzed per time point. (D) Effect of different induction levels of *lacZ* expression on the apparent transcription elongation rate, as calculated from FISH data. Error bars are standard deviations of at least three experiments. ** indicates a statistically significant difference (*P* < 0.01, two-sample *t* test). NS indicates a non-significant difference. (E) Distribution of Z5 mRNA numbers in a fluorescent spot inside cells at each time point for different IPTG concentrations. (F) Kinetics of the square root of LacZ activity when the promoter remained active (ON) or was turned off (OFF) 90 s after induction with 0.02 mM IPTG. AU, arbitrary units. Lines and shaded areas indicate the means and standard deviations of two-line fits on each time-course trace (five and three experiments for ON and OFF conditions, respectively). (G) Apparent transcription elongation rate of *lacZ* under conditions described in (F). Error bars show the standard deviations. NS indicates a non-significant difference. See also Figure S9 and Table S5.

A single RNAP was also largely insensitive to promoter activity, as we did not observe a significant delay in LacZ activity appearance when the *lac* promoter was turned off 90 s after induction with 0.02 mM IPTG (Figures 4F and S9). The apparent rate of transcription elongation was similar (*P* value = 0.42 from two-tailed t test) regardless of the promoter’s ON/OFF state (Figure 4G). Thus, the apparent slow-down in transcription elongation when the promoter is turned off is not a property of a single RNAP; instead, it is an emergent property of an RNAP group.

### Promoter shut-off promotes premature transcription termination

A significantly lower rate of transcription elongation often means longer or more frequent RNAP pauses and more efficient transcription termination (Fisher and Yanofsky, 1983; Guarente and Beckwith, 1978; Jin et al., 1992; Kotlajich et al., 2015; McDowell et al., 1994; Peters et al., 2011; Yanofsky and Horn, 1981). Thus, a potential functional consequence of RNAP stalling following the repression of an active promoter may be an increase in premature transcription termination. Time-course analysis of FISH data revealed that, under continuous induction, the Z5 and Z3 signals reached a similar plateau at steady state (Figure 5A), leading to a Z3/Z5 ratio close to 1 for various IPTG concentrations (Figure 5B). Since the degradation rates of the Z3 and Z5 regions were the same (with a mean lifetime of ~1.5 min, Figure S10), these results indicate that premature termination during *lacZ* transcription is negligible when the promoter remains active, as previously reported (Iyer et al., 2016). In contrast, when the promoter was shut off at 90 s, only ~50% of the RNAPs that transcribed the Z5 probe region reached the Z3 region (Figures 5C and 5D). Thus, a reduced elongation rate in response to a block in transcription initiation is associated with a significant increase in premature dissociation of the already-loaded RNAPs. For polycistronic genes, such as the *lac* operon, this premature transcription termination also suppresses the expression of downstream genes. In nature, where bacteria experience rapidly changing environments, this premature termination of transcription would be advantageous, as cells can more quickly stop the production of unneeded proteins when the inducing conditions disappear.

**Figure 5.**
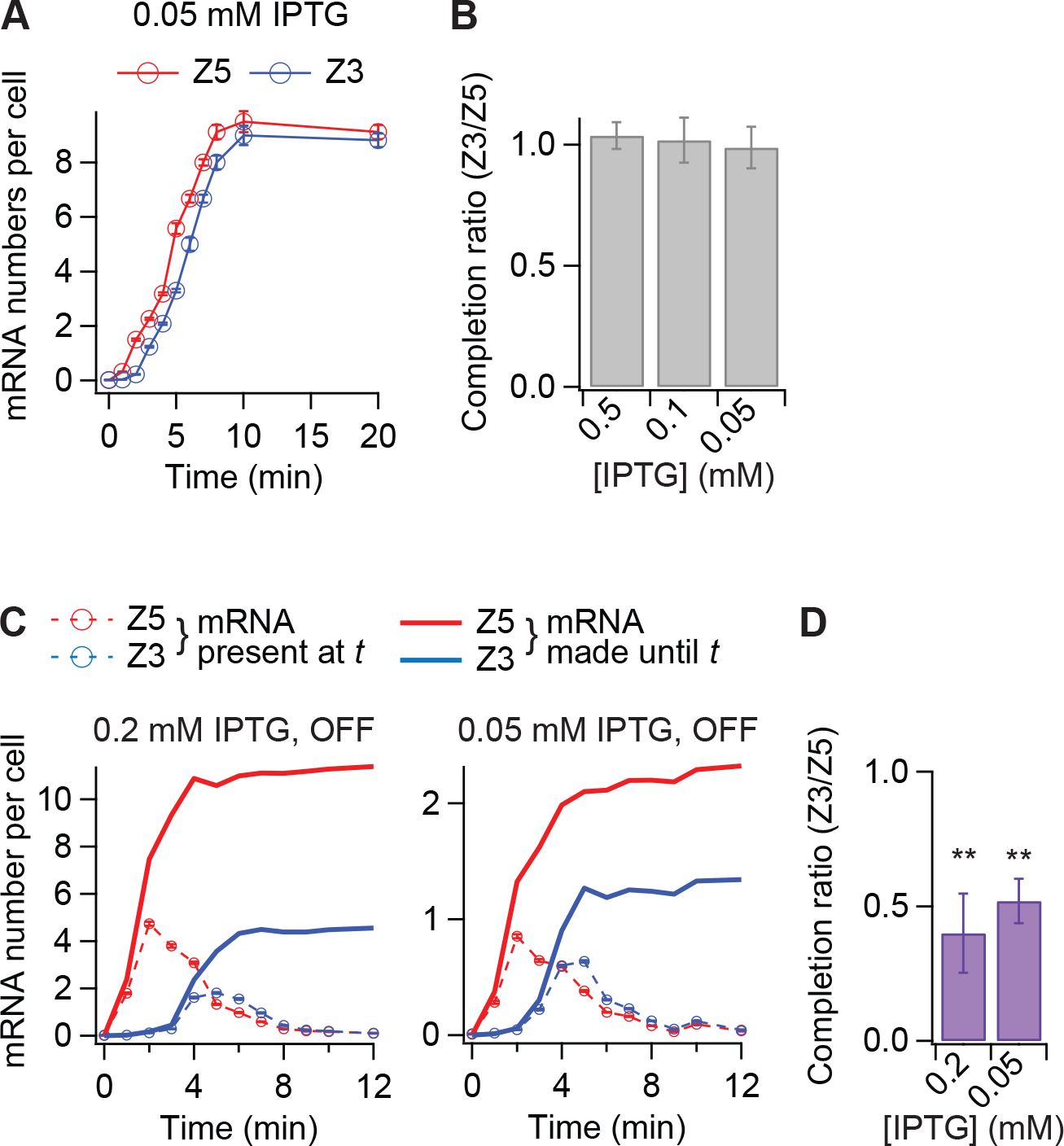
Premature dissociation of already-loaded RNAPs following promoter inactivation. We estimated the fraction of RNAPs that transcribe the Z5 region and also reach the Z3 region by examining the amount of Z5 and Z3 synthesis at the end of the time-course experiment. (A) Temporal change in the mean Z5 and Z3 mRNA numbers per cell under continuous induction of *lacZ* expression with 0.05 mM IPTG (promoter “ON”). Over 1500 cells were analyzed per time point. Error bars are bootstrapped standard errors of the mean. (B) Transcription completion ratio, i.e., ratio of RNAPs completing transcription in (A), calculated by dividing the Z3 plateau level by that of Z5. Over 7500 cells were analyzed for each IPTG concentration. Error bars are standard deviations of four, four, and five experiments for the 0.5, 0.1, and 0.05 mM IPTG conditions, respectively. (C) Accumulation of Z5 and Z3 mRNA numbers per cell when the promoter is turned off at *t* = 90 s. The total number of Z5 and Z3 mRNAs made until each time point (solid line) was calculated from their FISH signals (circles and dotted lines) using eq. 3 (see Methods). Over 2000 cells were analyzed per time point. Error bars are bootstrapped standard errors of the mean. (D) Transcription completion ratio, calculated from (C) by dividing the plateau level of Z3 by that of Z5. Error bars are standard deviations of four and six experiments for the 0.2 and 0.05 mM IPTG conditions, respectively. *** indicates a statistically significant difference to 1 (*P* < 0.001, one-sample *t* test). See also Figure S10.

### Expression from a gene can impact the transcription elongation rate of a divergently transcribed gene

Our proposed mechanism may also have implications for neighboring genes. It is already established that negative DNA supercoiling created during transcription can promote the local unwinding of DNA and facilitate transcription initiation of a neighboring gene if it is transcribed in the opposite direction (Dunaway and Ostrander, 1993; Meyer and Beslon, 2014; Naughton et al., 2013b; Opel and Hatfield, 2001; Rhee et al., 1999). Our model suggests that negative DNA supercoiling created by RNAP translocation on a gene may also reduce the speed of RNAPs on a neighboring divergent gene.

To test this prediction, we inserted *gfp*, driven by either a strong or a weak promoter, between *lacI* and *lacZ* on a plasmid in the Δ*lacZYA* strain (Figure 6A). Both promoters were derived from the *E. coli ompA* promoter, which we mutated to modulate its strength (Figures S11A and S11B). Without IPTG induction, basal *LacZ* activity was, as expected, higher when *gfp* was driven by the strong promoter compared to the weak promoter or the control template lacking *gfp* (Figure S11C). In addition, *gfp* expression from the strong promoter reduced the apparent transcription elongation rate of *lacZ* when its expression was induced with 0.2 mM IPTG (Figure 6B), consistent with our model prediction. Thus, an antagonistic dynamics can also emerge from RNAPs on separate genes.

**Figure 6.**
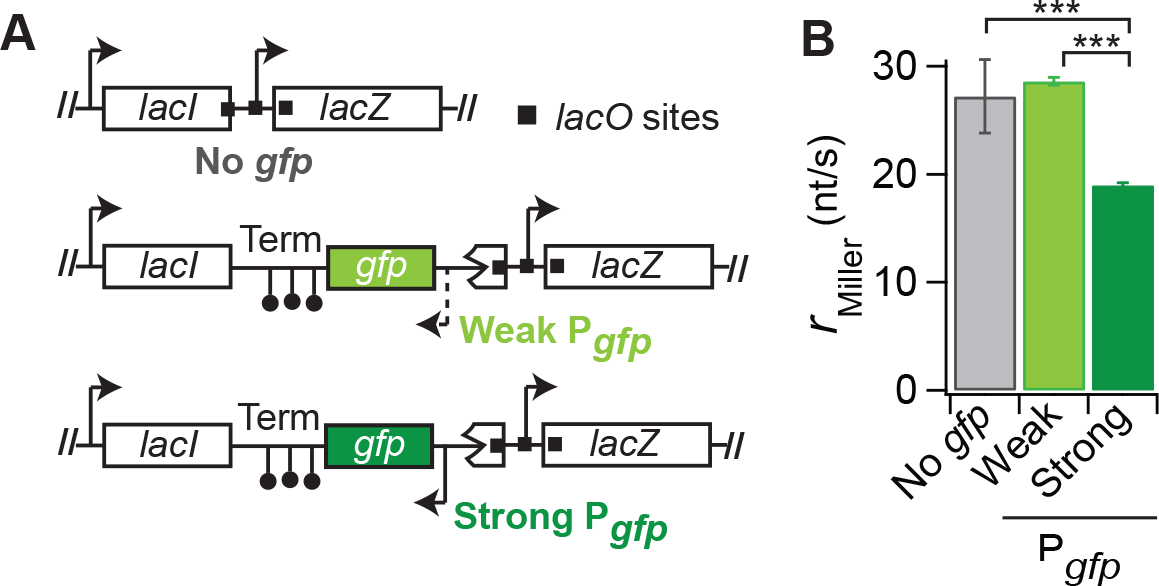
Effect of a divergently transcribed gene on *lacZ* transcription elongation. (A) Schematics of constructs used to test the effect of upstream divergent gene activity on transcription elongation of *lacZ* (not drawn to scale). (B) Apparent transcription elongation rate of *lacZ* for the different constructs, as measured by Miller assay under 0.2 mM IPTG induction (as in Figure 1C). The error bars show the standard deviations of three (no *gfp*), four (weak P_*gfp*_) and six (strong P_*gfp*_) experiments. *** indicates *P* < 0.001 (two-sample *t* test). See also Figure S11.

### Transcription-induced DNA supercoiling mediates RNAP group behaviors over long distances

Previous work has shown that when two RNAPs collide, the trailing RNAP can help the leading RNAP escape a pause site or overcome an obstacle, such as a DNA-binding protein or a nucleosome (Epshtein and Nudler, 2003; Epshtein et al., 2003; Jin et al., 2010; Saeki and Svejstrup, 2009). In this study, we show that co-transcribing RNAPs also display group behaviors over long (kilobase) distances (i.e., without collisions). These long-distance group behaviors emerge from two well-established properties of transcription on topologically constrained DNA: 1) the translocation of a single RNAP generates DNA supercoiling (Liu and Wang, 1987) and 2) DNA supercoiling impedes the motion of individual RNAPs (Chong et al., 2014; Ma et al., 2013; Rovinskiy et al., 2012). Our data suggest that these two properties, together with the ability of positive and negative DNA supercoils to diffuse rapidly over long distances (van Loenhout et al., 2012), can lead to both positive and negative effects among well-separated RNAPs.

Collectively, our data proposes the following model. When the promoter remains active, the presence of multiple RNAPs on the DNA template results in fluid RNAP translocation. DNA supercoils created by the translocation of each RNAP are rapidly cancelled out between RNAPs, relieving torsional stress on these RNAPs and leading to fast and processive translocation (Figure 3B). This long-distance assistance is not additive as the mechanism does not benefit from an increase in RNAP density. As a result, the elongation rate does not increase with the initiation rate as long as there are multiple RNAPs translocating on the same template (Figure 1). This long-distance assistance disappears when a single RNAP is transcribing or when an active promoter shuts off because torsional stress is no longer relieved by DNA supercoiling cancellation (Figure 3B). This results in slower elongation rates (Figures 2, 3A, and 4). In the case of promoter inactivation, the negative effect associated with the stalling of the promoter-proximal RNAP is quickly propagated to downstream RNAPs, as each of them benefits from the motion of the upstream RNAP for torsional stress relief.

Note that the *r* values for the promoter shut-off experiments (Figures 2B, 2D, and 3A, S5, S6, S7 and S8) underestimate the reduction in apparent elongation rate when the promoter becomes inactive. This is because the *r* values are calculated from the time of induction and therefore take into account not only the elongation rate after the promoter is shut off but also before it was shut off, i.e., when transcription elongation was fluid and faster. As discussed above (see text related to Figure 2), we estimate that it takes about three times longer for RNAPs to finish the last ~300 bp of *lacZ* transcription when the promoter is turned off at 90 s compared to when the promoter remains active. This implies that the average elongation rate is reduced from ~30 nt/s down to ~10 nt/s upon promoter inactivation, which is considerably lower than the average elongation rate of ~20 nt/s for a single RNAP (Figures 4B and 4D). In other words, RNAPs appear to translocate slower than a single RNAP when the promoter is turned off. How is this possible? We speculate that this is again linked to the ability of DNA supercoils to diffuse. RNAPs form bulky complexes with nascent transcripts and their associated ribosomes, and likely act as barriers to DNA supercoil diffusion (Leng et al., 2011). Therefore, the torsional stress experienced by RNAPs within a group after promoter inactivation may be higher than that experienced by a single RNAP because the DNA supercoils created between RNAPs are spatially confined compared to those created by a single RNAP. Furthermore, spatial confinement of DNA supercoiling may increase torsional stress due to the formation of a plectoneme (loop of helices twisted together), as the likelihood of plectoneme formation increases when DNA supercoiling occurs on shorter DNA segments (Brutzer et al., 2010).

The switch from collaborative to antagonistic group behavior following promoter inactivation is accompanied by a significant increase in premature termination (Figures 5C and 5D), presumably as a result of torsional stress and RNAP stalling. Prior to our work, the general assumption was that promoter inactivation in response to a change in intracellular or environmental conditions stops the loading of RNAPs, but does not affect the already loaded RNAPs. These RNAPs were assumed to continue transcription elongation normally, creating a wasteful delay between promoter inactivation and protein synthesis arrest. This would be analogous to stopping a car by taking the foot off the accelerator and not using the brake. However, our study shows that transcription from a group of RNAPs provides a built-in brake that more rapidly halts the production of proteins that are no longer needed.

Our data are also consistent with an emergent group function that can negatively impact RNAPs from divergently expressed gene pairs. If a gene is strongly expressed, negative DNA supercoils created by RNAP translocation can diffuse and impede the translocation of RNAPs on the neighboring divergent gene (Figure 6). Given the prevalence of divergent transcription in genomes (Wei et al., 2011), our result suggests another potential DNA supercoiling-dependent constraint on chromosomal gene arrangement during evolution (Meyer et al., 2018; Sobetzko, 2016). Our result also has implications for genetic engineering. Specifically, if fast transcription elongation is a desired property, one should avoid placing a pair of two strongly expressed genes in opposite directions.

Importantly, transcription-induced DNA supercoiling is a common feature of living cells across organisms (Giaever and Wang, 1988; Kouzine et al., 2014; Liu and Wang, 1987; Naughton et al., 2013a). Therefore, our findings may be broadly applicable, including to eukaryotic transcription.

## Supplemental Information

Supplemental Information includes eleven figures and five tables.

## Author contributions

S.K. and C.J.-W. designed the study. S.K. performed experiments and S.K., B.B., and C.J.-W. analyzed and discussed the data. S.K., B.B., and. I.I. provided resources. S.K. and C.J.-W. wrote the manuscript. All authors contributed to its editing.

## Acknowledgements

We thank Drs. Carol Gross, Jonathan Weissman, Matthew Larson, Hendrik Osadnik, and Jeff Hussmann for helpful discussions and technical help, Drs. Samuel Kou and Andrei Kuzminov for useful suggestions, Drs. Wayne Wade and Jeorg Bewersdorf for materials, and Dr. Andrew Goodman for allowing us to use his RT-PCR machine. We also thank Dr. Patricia Rosa and the members of the Jacobs-Wagner laboratory for discussions and critical reading of the manuscript. C.J.-W. is an Investigator of the Howard Hughes Medical Institute.

